# Knotted artifacts in predicted 3D RNA structures

**DOI:** 10.1101/2024.03.04.583268

**Authors:** Bartosz A. Gren, Maciej Antczak, Tomasz Zok, Joanna I. Sulkowska, Marta Szachniuk

## Abstract

Unlike proteins, RNAs deposited in the Protein Data Bank do not contain topological knots. Recently, admittedly, the first trefoil knot and some lasso-type conformations have been found in experimental RNA structures, but these are still exceptional cases. Meanwhile, algorithms predicting 3D RNA models have happened to form knotted structures not so rarely. Interestingly, machine learning-based predictors seem to be more prone to generate knotted RNA folds than traditional methods. A similar situation is observed for the entanglements of structural elements. In this paper, we analyze all models submitted to the CASP15 competition in the 3D RNA structure prediction category. We show what types of topological knots and structure element entanglements appear in the submitted models and highlight what methods are behind the generation of such conformations. We also study the structural aspect of susceptibility to entanglement. We suggest that predictors take care of an evaluation of RNA models to avoid publishing structures with artifacts, such as unusual entanglements, that result from hallucinations of predictive algorithms.

**Author summary:**

- 3D RNA structure prediction contests such as CASP and RNA-Puzzles lack measures for topology-wise evaluation of predicted models. Thus, predictors happen to submit potentially inappropriate conformations, for example, containing entanglements that are prediction artifacts.
- Automated identification of entanglements in 3D RNA structures is computationally hard. Distinguishing correct from incorrectly entangled conformations is not trivial and often requires expert knowledge.
- We analyzed 3D RNA models submitted to CASP15 and found that all entanglements in these models are artifacts.
- Compared to non-ML, machine learning-based methods are more prone to generating entanglements that are not present in natural RNAs.
- To increase the reliability of 3D RNA structure prediction, it is necessary to reject abnormally entangled structures in the modeling stage.

## Introduction

The birth of the third decade of this century brought a sudden surge of interest in modeling 3D RNA structures. The latter was, among other things, a by-product of the COVID-19 pandemic, whose main actor was the RNA virus, and intensive research on the development of RNA-based vaccines against COVID-19. The second major factor was the spectacular success brought about by the application of deep neural networks to model protein structures [1]. As a result, new methods have emerged to predict RNA structures, most of which use machine learning models in an end-to-end approach or at selected stages of the 3D folding process [2]. Many of them have undergone virgin benchmarking while competing in recent RNA-Puzzles and CASP15 initiatives, which aim to blindly evaluate predicted 3D RNA models and identify the best predictive tools. The results of both competitions show that none of the new methods has made a breakthrough in the quality and accuracy of the prediction of the 3D RNA structure so far. The latter are evaluated using various measures of distance (DI, GDT-TS, lDDT, MC☯, RMSD), similarity (INF, TM-score, LCS), and quality (Clash score) [3–7]. No measure can directly assess the accuracy of the 3D model topology and its compatibility with the target topology. Consequently, awareness of topological irregularities in 3D RNA predictions is negligible in the RNA community, and the predicted models happen to contain them. These anomalies include entanglements that are absent from known experimental structures. We can consider them locally, taking into account the secondary and tertiary structure of the molecule (we then speak of entanglements of structural elements), and globally, studying the spatial arrangement of the RNA backbone (we then analyze topological knots).

Entanglement of structural elements occurs when two elements of the RNA structure are in spatial conflict, that is, one of them punctures the other to form a lasso, interlace (known as a link in the knot theory) [8] or genus type (the genus trace represents how interconnected and densely packed the structure is in three dimensions) [9]. Loops, double-stranded fragments (consisting of dinucleotide steps), and single-stranded fragments can be involved in entanglements of this type. From the viewpoint of knot theory, lassos are not knots. Interlaces can be interpreted as Hopf links defined on loops traced by the phosphodiester and hydrogen bonds; the former contribute to the formation of the nucleotide chain and the latter to base pairs. The formation of many types of entanglements of structure elements, like some interlaces (loop & dinucleotide step, dinucleotide step & dinucleotide step) and deep lassos, contradicts the RNA folding hierarchy and is therefore hardly possible. We do not find them in high-resolution experimentally determined RNAs [10]. They occur in structures modeled *in silico*, in which case they should be regarded as artifacts of computational procedures.

The formation of topological knots in RNA structures has been studied by a few research teams so far. The authors of [11], although they came across knotted 16S rRNA domains, argued about the low likelihood of topological knots in large native RNAs and suggested an examination of experimental RNA models for knotting before their publication. Micheletti *et al*. [12] found three knotted rRNAs solved by cryo-EM, but due to the absence of knots in high-resolution structures, concluded on some thermodynamic or kinetic mechanisms that minimize the entanglement of biologically viable structural RNAs. Yet a little later, the same group suggested that the properties of some of the predicted RNA secondary structures indicate the potential to form knots [13]. Only recently, a trefoil (3_1_), the first non-trivial topological knot, was identified in high-resolution experimental RNA structure [14]. However, this is not sufficient to infer the formation of knots in RNA structures and their possible influence on the function of the molecule, as we can for proteins [15].

In this work, we have analyzed all 3D RNA models predicted in the recent CASP competition. Using existing computational tools, RNAspider [10] and Topoly [16], we have scanned these predictions for entanglements of structural elements and topological knots. We have studied the structural and methodological aspects of susceptibility to entanglement generation in RNA models. All entanglements found are artifacts of the modeling procedures. Methods using deep learning entangle RNA chains more often than non-ML algorithms and generate quite complex topological knots. We believe that predictive methods should automatically reject models with invalid chain entanglements. This would improve the reliability and quality of the prediction of the 3D RNA structure.

## Materials and methods

### Benchmark data

To identify, count, and classify knots and entanglements of structural elements in the 3D RNA structure models predicted in CASP15, we downloaded the data from the competition website in September 2023. The CASP15 resources are available at https://predictioncenter.org/download_area/CASP15/predictions/RNA/ [17]). Data were grouped by 12 RNA targets: R1107, R1108, R1116, R1117, R1126, R1128, R1136, R1138, R1149, R1156, R1189 and R1190. The data set contained a total of 62 reference structures (for some targets, there was more than one structure) and 1,660 models computationally predicted by 41 modeling groups.

### Methods used for entanglements of structure elements

The RNAspider web server was applied to identify and classify entanglements of structure elements in 3D RNA structures [10]. The system detects lassos and interlaces and assigns them to nine subclasses (Fig. 1A,B) – L&L, L&D, D&D, L(S), L(D), L(L), D(S), D(D), and D(L) – based on the types of elements involved, i.e., loops, dinucleotide steps (both are closed structural elements), and single-stranded fragments (open element). It was run with the default settings of the advanced parameters. We then calculated the depth of each identified lasso formed from the loop L(*), where * stands for S, D, or L. This was done using an additional script, as RNAspider does not cover such functionality. The script was fed with information on the intersection points that RNAspider provided for every closed structure element. The intersection point is determined by 3D coordinates, where the backbone or hydrogen bond in a canonical base pair punctures a surface that covers a closed element. We considered two cases. In the case of L(S), when a single strand is taken to lasso by the loop, we computed the depth as the minimum number of its nucleotides towards the 5’ and 3’ ends from the intersection point. In the second case, when a closed element is lassoed (L(L), L(D)) or when a single strand punctures the loop twice (L(S.)), the lasso is characterized by two intersection points. Therefore, we computed the depth as the number of nucleotides between the intersection points. Knowing the depth, we divided the lassos created by the loops into shallow (depth *≤* 5 nts) and deep (depth *>* 5 nts). Following our experience with molecular dynamics, we hypothesize that shallow lassos L(*) can spontaneously disentangle during structure folding and therefore may not be treated as structure anomalies even though no lassos occur in the reference structures. On the contrary, we consider interlaces, D(*) lassos, and deep L(*) lassos as modeling artifacts.

**Fig 1.**
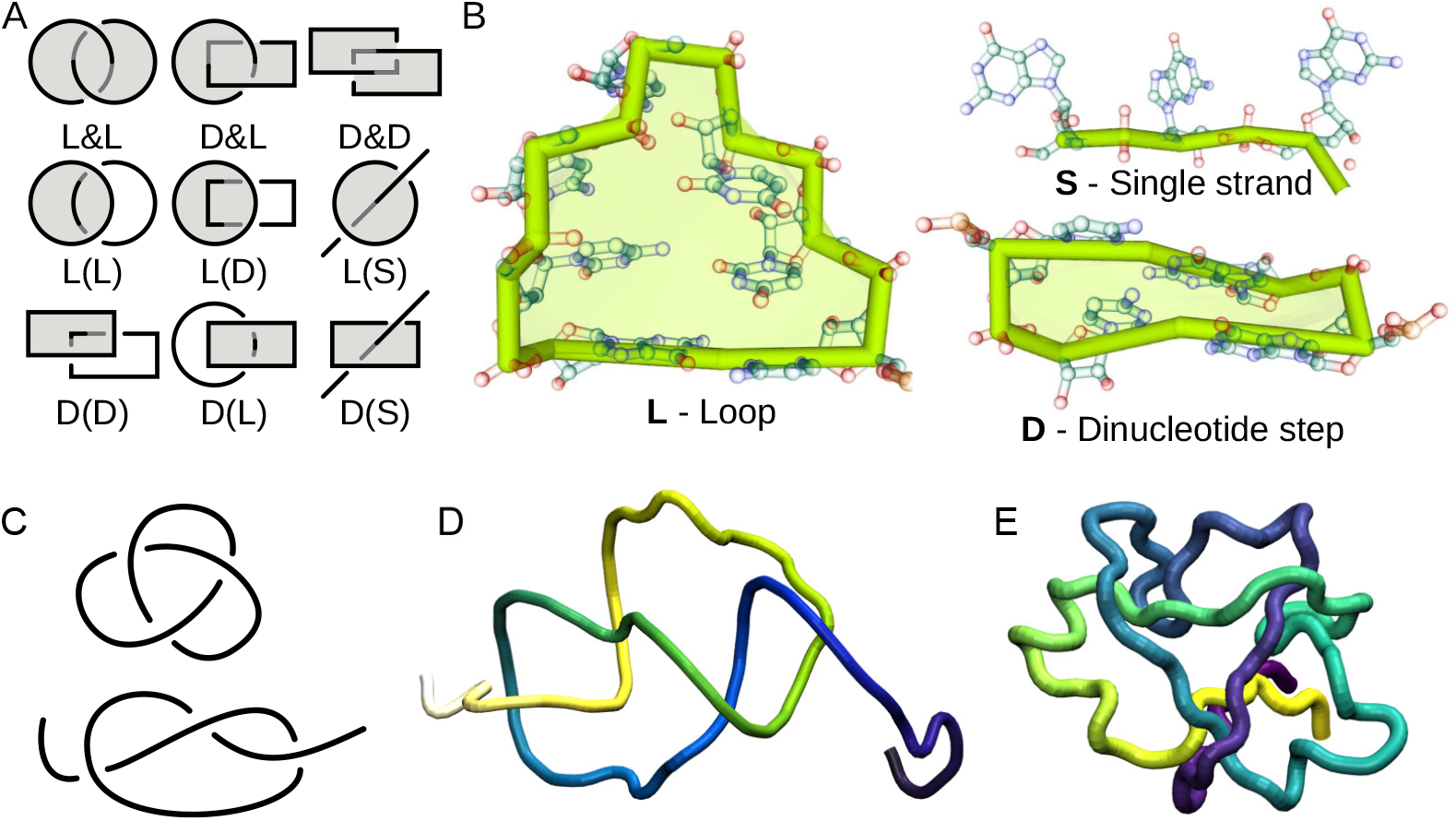
Types of entanglements. (A) Schematic drawings of interlaces (D&D, D&L, L&L) and lassos (D(D), D(L), D(S), L(D), L(L), L(S)), (B) example structural elements L, D, and S, (C) diagrams of closed and opened trefoil knot, and two molecules with trefoils - (D) sRNA RydC (PDB ID: 4V2S:G [18]) and (E) PHD finger-like domain-containing protein 5A (PDB ID: 5ZYA:C [19]).

### Methods used for topological knots

Topological knots were detected and identified using the Topoly Python package [16]. This collection of scripts offers features to study the topology of polymers and a generation of artificial, random polymer chains of a given topology. Here, the coordinates of the sugar-phosphate backbone atoms (P, O5*′*, C5^*′*^, C4^*′*^, C3^*′*^, O3^*′*^) were extracted from the 3D structure data. To identify knots, the Alexander polynomial [20] was calculated using the `topoly.alexander()` function, which before calculations closes the chain randomly by projecting RNA endpoints on the big sphere around the molecule and connecting them (two-point probabilistic closure, 200 closures, explained more in [21, 22]). We treated the structure as knotted if *<* 50% of the closures were unknots. At first, Topoly identified 84 knotted RNAs. They were visually inspected to confirm entanglements. For some of them, a direct closure seemed to make more sense. We discarded 7 of these models since they were unknots after direct closure. 67 predictions failed to process by Topoly due to non-unique atom coordinates (47 models) or too tangled structure (20 models). Structures in the latter subset were visually inspected and confirmed to be too densely packed to be correct. We assigned them to a category named TTC (too tangled to check) and did not include them in further analysis, unlike trefoils (Fig. 1C) and complex knots with known classification.

## Results

In this work, we analyzed entanglements in 3D RNA models submitted to CASP15. We took into account the entanglements of structural elements and topological knots. It is important to emphasize that none of the reference structures, of which there were 62 models, was entangled from the point of view of the structure elements or topological knots. In contrast, of the 1,660 predicted models, 162 models have either entanglements of structure elements or topological knots, 83 models have only entanglements of structural elements, 34 models have only topological knots, and 43 models have both. Fig. 2A shows a Venn diagram detailing the number of predicted models that contain given types of entanglements. Note that the existence of topological knots in the 3D RNA model does not equate to entanglement of the structural elements of the model and vice versa. In 126 entangled models, we encountered 173 interlaces and 300 lassos (Fig. 2B). The latter group included 67 deep and 33 shallow conformations of the L(*) type. Note that among the lassos, the L(S) and L(D) types are most abundantly represented in the RNA predictions. Intuition dictates that this is entirely reasonable since forming a lasso around a single or double helix should be relatively easy. In contrast, pushing a non-single-stranded fragment of a structure through a dinucleotide step or a loop through a loop seems quite difficult. Therefore, these types of entanglements are relatively rare in the models. In 77 RNA models with topological knots, we found a predominance of trefoils - they account for more than half of all knots with assigned classes excluding the TTC subset (Fig. 2C). TTC, the second large subgroup, is made up of 20 models whose knotting is too complex to classify correctly.

**Fig 2.**
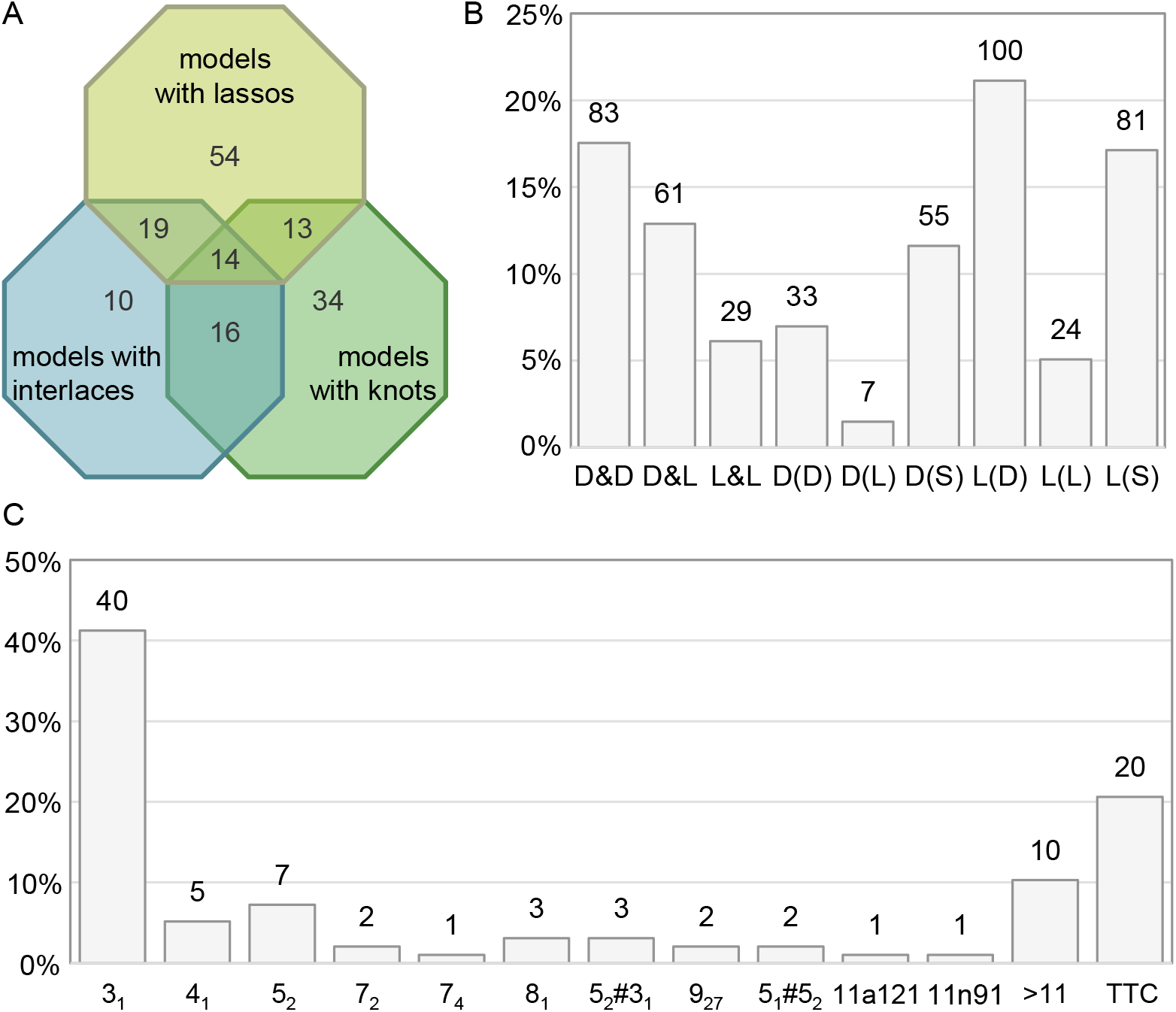
(A) Models with various entanglement types in numbers. (B) Entanglements of structure elements and (C) topological knots in RNA predictions by type. Column labels in (B) and (C) show the total number of entanglements of a given type across all predictions.

### Target-focused analysis

In this part of our experiment, we focused on the structural aspect of the structure entanglement problem. We asked whether, among the RNA targets in CASP15, we could distinguish structures that show a greater /lower susceptibility to entanglement during computer modeling. Answering this question required analyzing the predicted models clustered by target (see Fig. 3 and Table 1).

**Table 1.**
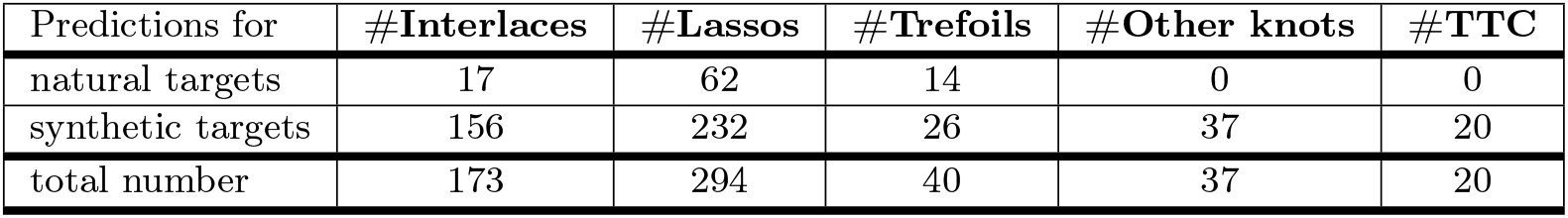
Types of entanglements in RNA predictions by target.

**Fig 3.**
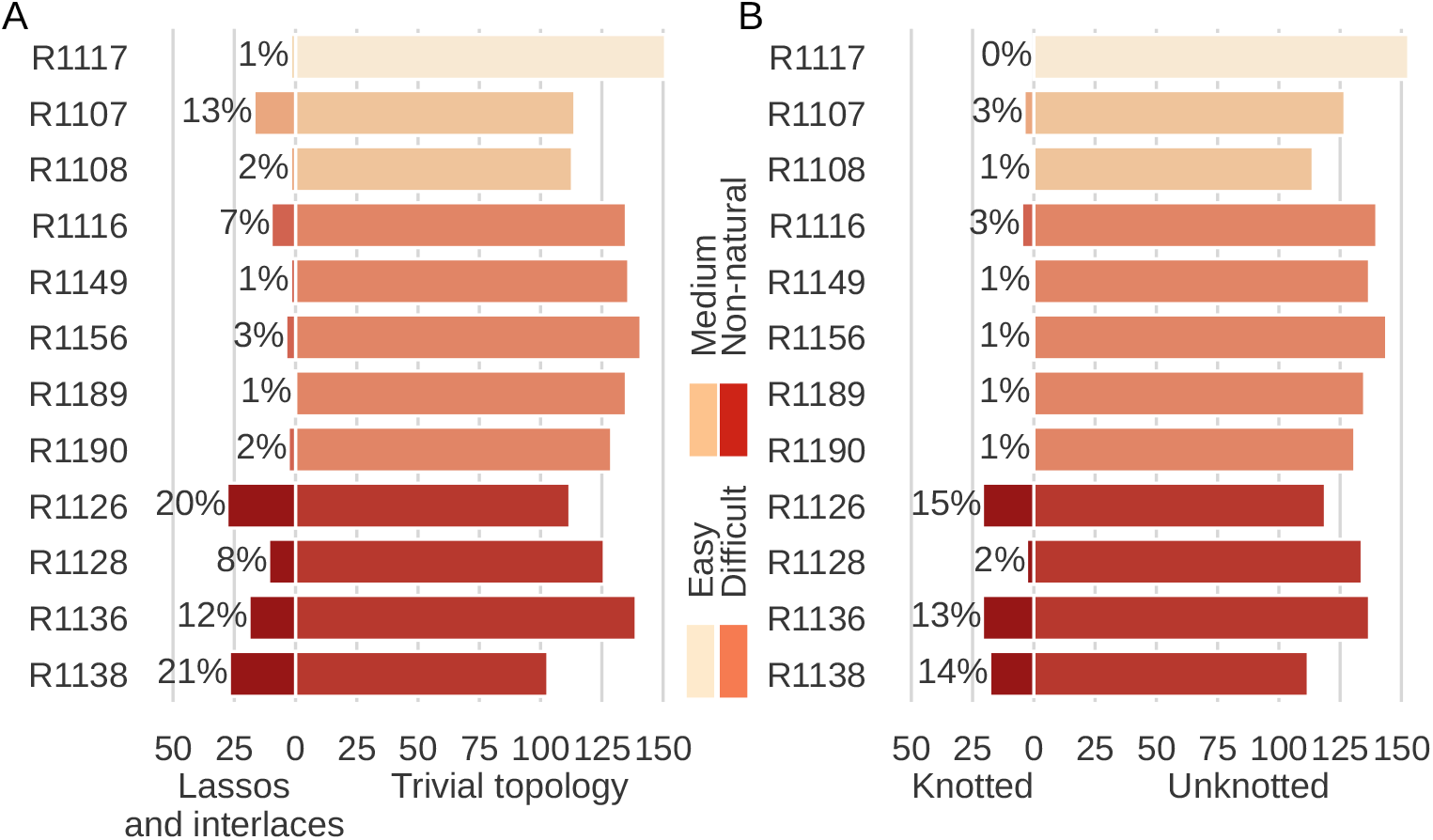
Distribution of entangled 3D RNA structure predictions by target. Target structures are grouped by difficulty [7]. Results are displayed separately for (A) entanglements of structure elements (lassos and interlaces) and (B) topological knots.

The target structures were divided by the CASP assessors into natural (8 targets) and synthetic (4 targets). The former cluster distinguished between easy (R1117), medium (R1107, R1108), and difficult (R1116, R1149, R1156, R1189, R1190) targets. Non-natural ones included R1126, R1128, R1136, and R1138. In general, 33% (41) of the entangled structures are predictions of natural RNA targets, and 67% (85) are models from the non-natural cluster.

Based on the analysis of entanglements of structural elements (Fig. 3A), we can say that the probability of entangled predictions for natural RNAs equals 0.03, while for non-natural targets it is 0.15, which is 5x higher. If we look at the sets of predictions per target, we can see that the entangled structures represent 8-20% for the synthetic targets. In contrast, in the sets of models for natural targets, the percentage of entangled predictions is 0.74-2.76%. The exceptions here are R1107 (12.98%) and R1116 (6.90%), the former of which is a moderately difficult structure with a pseudoknot, and the latter is classified as difficult.

The diagram prepared for the topological knots (Fig. 3B) has similar characteristics as in the case of entangled structure elements. The highest number of knotted models is found for the targets for which we observe the most entanglements of structural elements. The study reveals that the knotting probability is 0.01 for natural structures and 0.12 (10x higher) for non-natural ones. All knotted predictions of natural structures are trefoils (3_1_, simplest non-trivial knot). For non-natural structures, only more complex knots appear, moreover, they make up the majority of knotted structures.

Some predictions contain more than one entanglement. Table 1 presents a distribution of various types of entanglements across predictions for natural and synthetic targets. For simplicity, topological knots were divided into trefoils (the simplest non-trivial knot) and other knots (more complicated ones). The rightmost column represents unclassified knots from non-physically dense structures (TTC - too tangled to check knot type).

Finally, let us add that the largest number of entangled 3D models (both from the point of view of topological knots and entangled structure elements) was identified in the predictions for the three largest targets, R1138 (720 nts), R1136 (374 nts), and R1126 (363 nts), all of which are synthetic. However, note that, as shown in [8], no simple relationship has yet been observed between the entanglements of structural elements and the size of the structure. Given that topological knots more complex than trefoil are found only in the predictions of synthetic targets, it seems that the large number of entanglements and their complexity are a result of the specificity of the non-natural target rather than the structure size.

### Method-focused analysis

Next, we analyzed 3D RNA models by prediction group and examined the structure entanglement problem from the point of view of the method. We checked how many non-trivial topologies, distinguishing between topological knots, lassos, and interlaces, appear in the models generated by a given prediction method (Fig. 4). Of the 41 groups, 16 submitted only unentangled models (a total of 510 predictions, which makes 30% of all predictions). The remaining 25 groups predicted 1,150 models, of which 126 (10%) include entanglements of structure elements, and 78 (6%) are knotted from the topological point of view.

**Fig 4.**
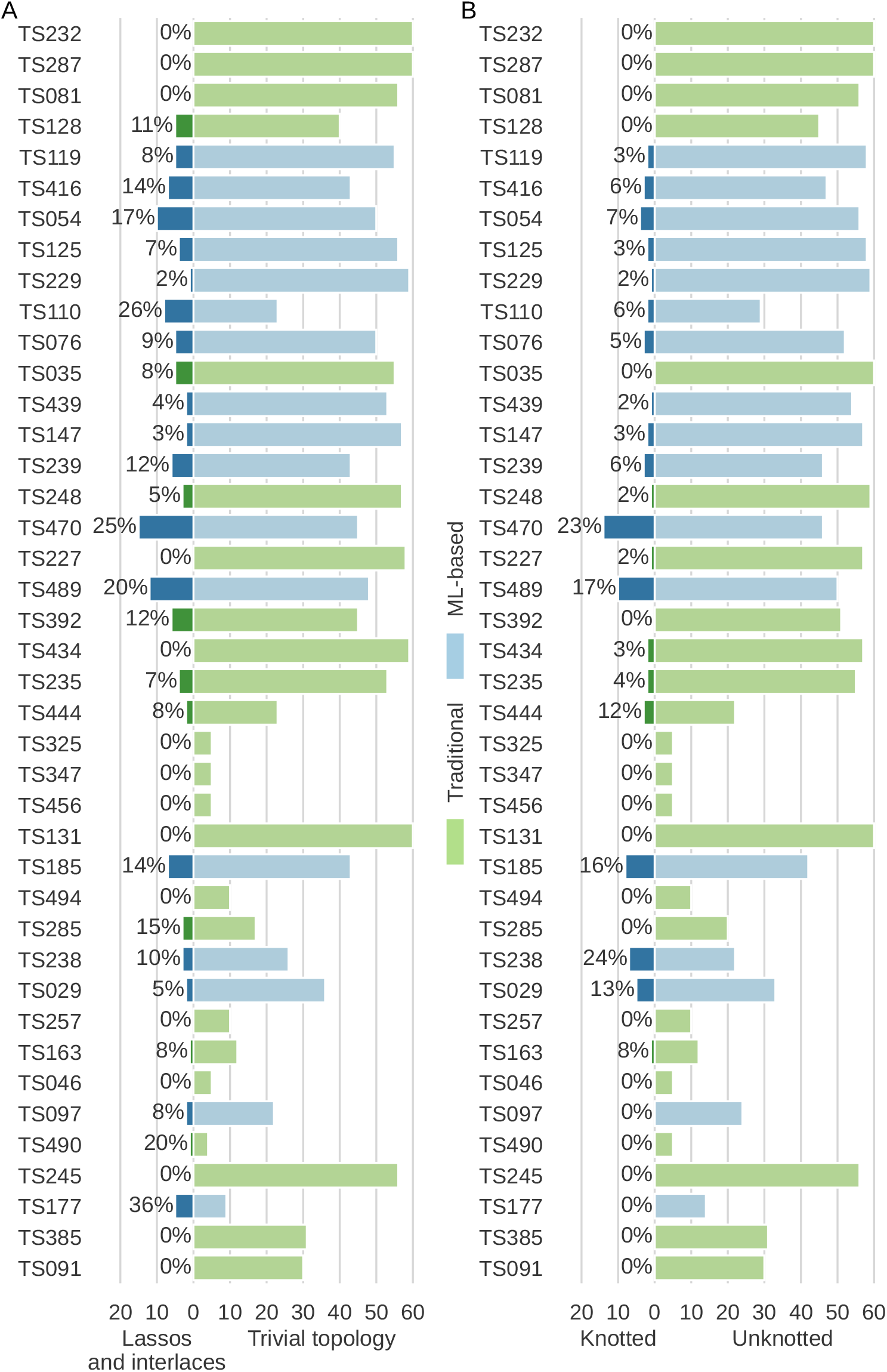
Distribution of entangled 3D RNA structure predictions by method. Modeling groups (classified as traditional or ML-based) are listed due to their ranking in CASP15 with the best one at the top. Results are shown separately for (A) entanglements of structure elements (lassos and interlaces) and (B) topological knots.

Among 126 RNA models with entangled structure elements, 101 (80%) were predicted using machine learning-based methods, and the remaining 25 (20%) by traditional approaches. This means that ML methods are 4x more prone to generate models containing entanglements of structural elements than traditional algorithms. A similar trend is observed when analyzing topological knots. Among 77 knotted models, 67 (87%) were predicted by machine learning-based methods, and the remaining 10 (13%) by traditional approaches. Thus, the susceptibility of ML methods to generate knotted models is *∼* 7x greater than that of non-ML approaches. ML methods predicted *∼* 6x more trefoils (34 vs 6) and *∼* 8x more knots which are more complicated than a trefoil. Moreover, all 20 structures, which were too tangled to check their knot type (TTC), were also predicted by ML methods. The distribution of the entanglement types is presented in Table 2.

**Table 2.** Types of entanglements in RNA predictions by method.

| Models predicted | #Interlaces | #Lassos | #Trefoils | #Other knots | #TTC |
| --- | --- | --- | --- | --- | --- |
| by traditional methods | 62 | 103 | 6 | 4 | 0 |
| by ML-based methods | 111 | 191 | 34 | 33 | 20 |
| by servers | 47 | 66 | 7 | 10 | 8 |
| by human predictors | 126 | 228 | 33 | 27 | 12 |
| in total | 173 | 294 | 40 | 37 | 20 |

Finally, recall that CASP accepts submissions from two categories of participants, web servers, and human groups. Predictions from the former category are fully automatic and must be submitted within 72 hours of publishing the target sequence. Human groups have 3 weeks to make predictions and can utilize any method to support the modeling process, including laboratory experiments to refine their models. With this in mind, we checked whether topological knots and entanglements of structure elements are more often in the web server than in human predictions. Among the 41 participants, there were 9 web servers. They submitted a total of 423 models, including 29 (0.07%) with entanglements. 32 human groups predicted 1,237 models, of which 97 (0.08%) were entangled. The number of entanglements of each type in the models submitted by both categories of participants is shown in Table 2. Based on these data, the thesis that automated predictions are more likely to get entangled than those submitted by experts cannot be confirmed.

### Example predictions with artifacts

First, let us present the 3D RNA prediction that contains a lasso and was generated using the machine learning-based method. The R1107TS416_2 model was submitted by the TS416 group (AIchemy_RNA). It targeted the natural 69-nt-long RNA structure of the human CPEB3 HDV-like ribozyme (PDB ID: 7☯R4) [23] (target ID: R1107). The native structure contains a pseudoknot formed between the dangling 5’-end (residues 1-6) and the three-way junction (residues 31-36). The R1107TS416_2 model is not an ideal reconstruction of the target either from the point of view of secondary structure (INF_*all*_ = 0.77) or the 3D topology (RMSD = 7.81Å, TM-score = 0.392). The model contains the L(S)-type lasso formed between the 26-nt-long hairpin closed by a pseudoknotted base pair 0:6-0:31 and a 34-nt-long single strand (0:36-0:69) (Fig. 5A). Both of these structure elements exist also in the native structure, although in the latter a single strand bypasses the loop. In the predicted model, the intersection of the area inside the loop is between residues 0:A62 and 0:U63 of the puncture strand. The single strand passes quite a distance from the chain that forms the loop. Therefore, we did not observe clashed atoms in this part of the structure; in general, the Clash score is low and equals 14.02. According to the threshold that we adopted for shallow lassos, the entanglement found belongs to the deep category; its depth is equal to 6 nucleotides.

**Fig 5.**
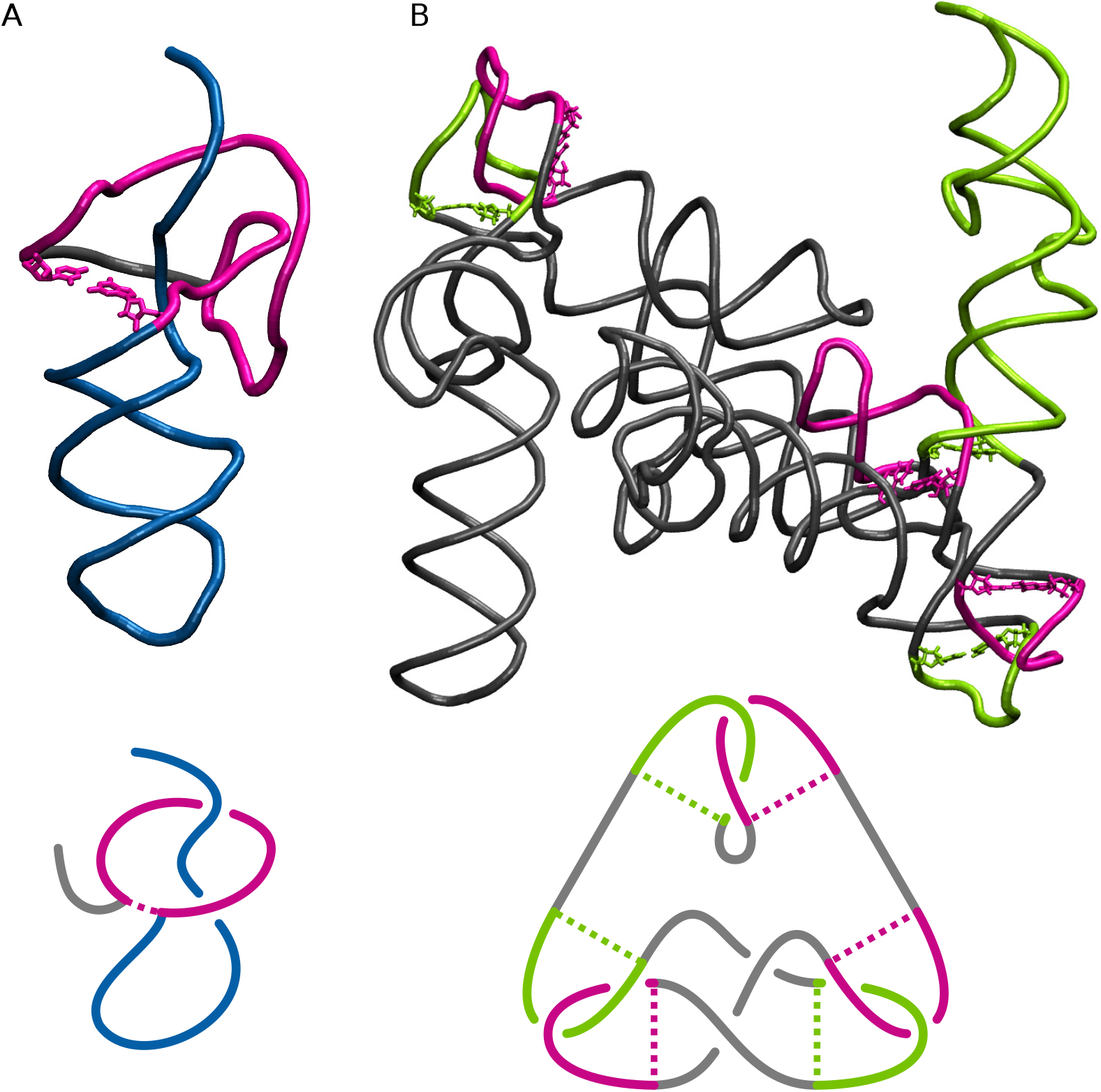
Two models with entangled structural elements predicted by ML-based methods and diagrams showing included entanglements. (A) The R1107TS416_2 model with a lasso. (B) The R1136TS110_4 model with highlighted three interlaces forming the 3_1_ knot. Hydrogen bonds closing an entangled loop are marked with dotted lines.

The other example, R1136TS110_4 model, was submitted by the TS110 group (DF_RNA). It is an ML-driven prediction of the synthetic construct, which is the 3D structure of a brocolli-pepper aptamer FRET tile in the ligand-bound state (PDB ID: 7ZJ4) [24] (target ID: R1136). The reference structure consists of 374 nucleotides and has a non-trivial topology with kissing-loop interactions between two hairpins (0:50-0:60+0:142-0:152; 0:75-0:85+0:121-0:131). The predicted model’s secondary structure is predicted quite well (INF_*all*_ = 0.82), while the 3D fold clearly deviates from the native (RMSD = 44.35Å, TM-score = 0.304). Clash score = 49.78. The model contains six entanglements of structural elements – two D(D), two D&D, one D&L, and one D(L) (Fig. 5B), formed between two 10-nt-long hairpin loops (0:142-0:152; 0:50-0:60) and seven dinucleotide steps (0:119-0:120+0:132-0:133; 0:159-0:160+0:216-0:217; 0:160-0:161+0:215-0:216; 0:335-0:336+0:345-0:346; 0:49-0:50+0:60-0:61; 0:336-0:337+0:344-0:345; 0:293-0:294+0:299-0:300). They are not only artifacts of modeling but also incorrect conformations hardly possible to form while RNA folding.

The next example presents different topological knots in the 3D RNA models predicted for the largest RNA target of CASP15, that is, R1138 (720 nts). The native structure (synthetic construct) is a young conformer of a 6-helix bundle of RNA with clasp (PDB IDs: 7PTK, 7PTL) [25]. As shown in Fig. 6A, it does not contain topological knots. Model R1138TS227_2 (Fig. 6B) was generated by a traditional (non-ML) approach used by the TS227 group (GinobiFold). RMSD = 48.23Å and TM-score = 0.204 indicate a significant deviation of its 3D topology from the target structure, while the secondary structure is quite well-reproduced (INF_*all*_ = 0.82).

**Fig 6.**
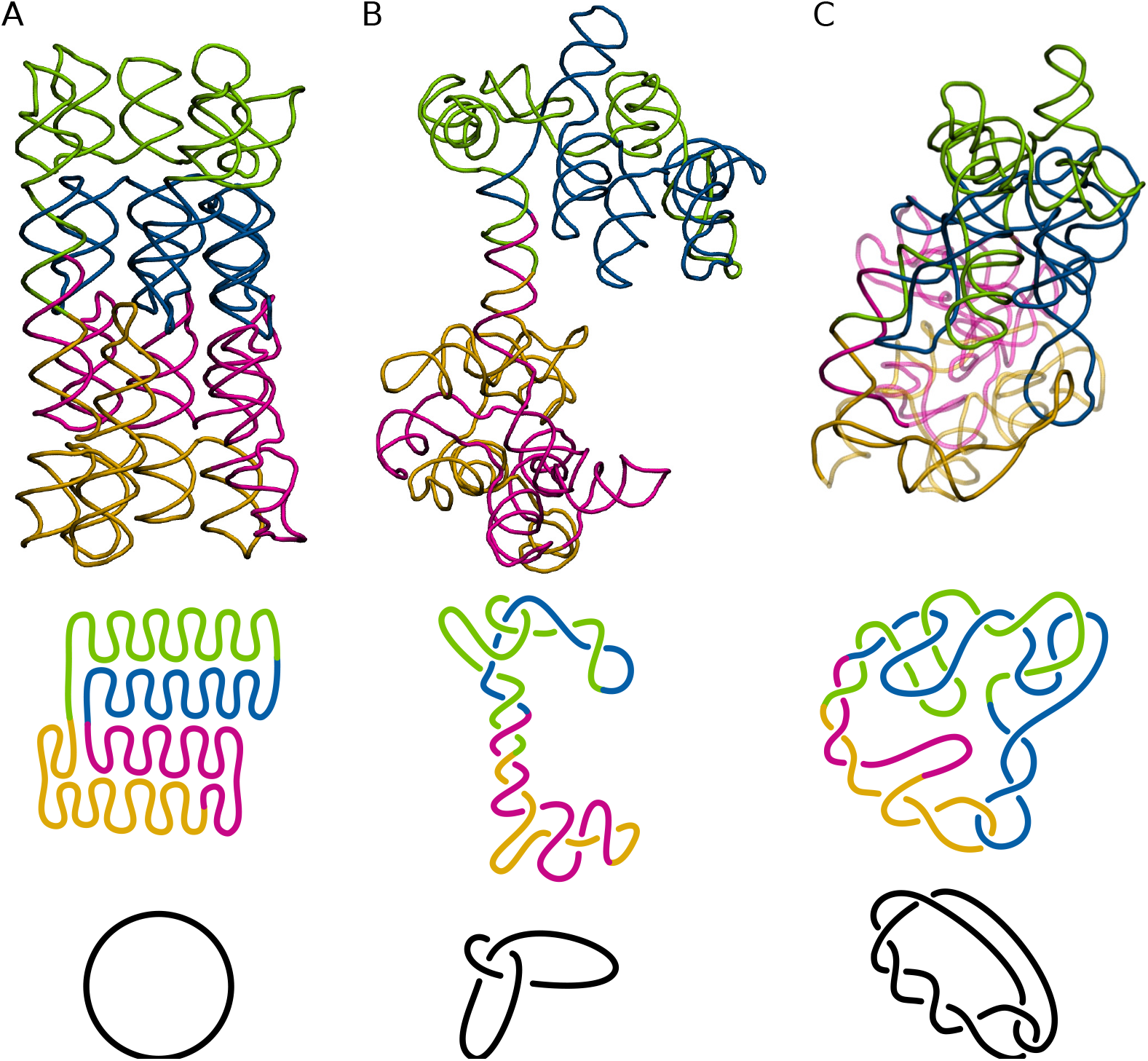
3D models predicted for the same target containing various topological knots and schematics of the latter. (A) Target structure (R1138). (B) The R1138TS227_2 model generated by traditional method. (C) The R1138TS054_3 model predicted by ML-based method.

Interestingly, clashes are almost non-existent (Clash score = 0.30). This model is knotted and forms a trefoil (3_1_), the simplest non-trivial knot. On the other hand, the R1138TS054_3 model was predicted using the machine learning-based method by the TS054 group (UltraFold). Its ratings are similar to the previous example (RMSD = 39.73Å, TM-score = 0.186, INF_*all*_ = 0.85, and Clash score = 4.81), however, this structure forms a more complicated 7_2_ knot. This clearly shows that existing evaluation measures are not correlated with the complexity of the structure’s entanglement.

## Conclusion

An analysis of the 1,660 3D RNA models predicted within CASP15 showed that predictive methods using machine learning are four times more likely than traditional methods to generate structures with entanglements, which are artifacts of the computational process. We hypothesize that the generation of entanglements may be due to the algorithms’ prioritization of more compact structures. Predictions submitted by web servers and by human groups contain the same percentage of entangled models. The implication is that predictors do not use any, automatic or non-automatic, verification of their models for topological anomalies. The characteristics of the modeled structure also appear to affect the probability of entanglement. In the set of all predictions, the largest number of entangled models was generated for large targets of synthetic RNA molecules. The probability of entanglement in this subset was significantly higher than for natural structures. We believe that enriching prediction methods with procedures to validate the topology would increase the accuracy of 3D RNA structure prediction.

## Acknowledgments

This work was supported by the National Science Centre, Poland (2018/31/B/NZ1/04016 and 2021/43/I/NZ1/03341 to JIS). MA, TZ, and MS acknowledge the support of the Poznan University of Technology and the Institute of Bioorganic Chemistry, PAS (statutory funds).

