## Supplemental Table 1 for "Knotted artifacts in predicted 3D RNA structures"

**S1 Table. Entangled 3D RNA models predicted in CASP15.**

| No | Model | Human/ Web server | ML/ nonML | Interlaces | Lassos | | Knots |
|---|---|---|---|---|---|---|---|
| | | | | | Shallow | Deep (>5) | |
| 1 | R1107TS054_1 | Human | ML | | | L(S) | $0_1$ |
| 2 | R1107TS054_3 | Human | ML | | | L(S) | $0_1$ |
| 3 | R1107TS054_4 | Human | ML | | | L(S) | $0_1$ |
| 4 | R1107TS054_5 | Human | ML | | | L(S) | $0_1$ |
| 5 | R1107TS119_2 | Human | ML | | L(L) | L(D) | $0_1$ |
| 6 | R1107TS119_3 | Human | ML | | | L(S) | $0_1$ |
| 7 | R1107TS125_1 | Web server | ML | | | L(S) | $0_1$ |
| 8 | R1107TS125_2 | Web server | ML | | | L(S) | $0_1$ |
| 9 | R1107TS128_2 | Human | nonML | | L(L) | L(D) | $0_1$ |
| 10 | R1107TS163_1 | Web server | nonML | | L(S) | L(S) | $0_1$ |
| 11 | R1107TS392_1 | Human | nonML | | | L(S) | $0_1$ |
| 12 | R1107TS392_2 | Human | nonML | | | L(S) | $0_1$ |
| 13 | R1107TS392_3 | Human | nonML | | | L(S) | $0_1$ |
| 14 | R1107TS392_4 | Human | nonML | | | L(S) | $0_1$ |
| 15 | R1107TS392_5 | Human | nonML | | | L(S) | $0_1$ |
| 16 | R1107TS416_2 | Human | ML | | | L(S) | $0_1$ |
| 17 | R1107TS416_3 | Human | ML | | | L(S) | $0_1$ |
| 18 | R1108TS119_5 | Human | ML | | | L(S) | $0_1$ |
| 19 | R1108TS128_2 | Human | nonML | | L(L) | L(D) | $0_1$ |
| 20 | R1116TS029_3 | Human | ML | | | L(S) | $0_1$ |
| 21 | R1116TS035_3 | Web server | nonML | L&L | | L(D) | $0_1$ |
| 22 | R1116TS035_4 | Web server | nonML | L&L | | L(D) | $0_1$ |
| 23 | R1116TS035_5 | Web server | nonML | L&L | | L(D) | $0_1$ |
| 24 | R1116TS177_1 | Human | ML | | | L(S) | $0_1$ |
| 25 | R1116TS248_3 | Human | nonML | | | L(S) | $0_1$ |
| 26 | R1116TS285_4 | Human | nonML | | L(L) | L(D) | $0_1$ |
| 27 | R1116TS470_1 | Human | ML | D&D | | D(D) | $0_1$ |
| 28 | R1116TS470_3 | Human | ML | D&D,D&L | | L(D) | $0_1$ |
| 29 | R1116TS470_5 | Human | ML | D&D | | | $0_1$ |
| 30 | R1117TS097_1 | Human | ML | | L(S) | | $0_1$ |
| 31 | R1117TS125_4 | Web server | ML | | L(S) | | $0_1$ |
| 32 | R1126TS029_1 | Human | ML | | | | $3_1$ |
| 33 | R1126TS029_2 | Human | ML | | | | $5_2$ |
| 34 | R1126TS035_1 | Web server | nonML | | L(L) | | $0_1$ |
| 35 | R1126TS054_1 | Human | ML | | | L(D) | $0_1$ |
| 36 | R1126TS076_2 | Human | ML | L&L | | | $3_1$ |
| 37 | R1126TS110_1 | Human | ML | D&D | | | $0_1$ |
| 38 | R1126TS110_2 | Human | ML | D&D | | | $0_1$ |
| 39 | R1126TS110_3 | Human | ML | D&D | | | $0_1$ |
| 40 | R1126TS128_2 | Human | nonML | | | L(D) | $0_1$ |
| 41 | R1126TS147_5 | Human | ML | | | L(D) | $0_1$ |
| 42 | R1126TS177_1 | Human | ML | | | L(D),L(S) | $0_1$ |
| 43 | R1126TS185_1 | Human | ML | | | L(D) | $0_1$ |
| 44 | R1126TS185_2 | Human | ML | | | L(D) | $0_1$ |
| 45 | R1126TS185_4 | Human | ML | D&L | | L(S) | $4_1$ |
| 46 | R1126TS185_5 | Human | ML | L&L | | L(D),L(S) | $0_1$ |
| 47 | R1126TS238_1 | Human | ML | | | | 12a380 or 2n307 or $7_2\#5_2$ |
| 48 | R1126TS238_2 | Human | ML | | | | >12 |

| No | Model | Human/ Web server | ML/ nonML | Interlaces | Lassos | | Knots |
|---|---|---|---|---|---|---|---|
| | | | | | Shallow | Deep (>5) | |
| 49 | R1126TS238_3 | Human | ML | | | | >12 |
| 50 | R1126TS239_1 | Web server | ML | L&L | | | $3_1$ |
| 51 | R1126TS239_5 | Web server | ML | | | D(S) | $0_1$ |
| 52 | R1126TS248_1 | Human | nonML | | L(L) | | $0_1$ |
| 53 | R1126TS416_4 | Human | ML | | | | $3_1$ |
| 54 | R1126TS416_5 | Human | ML | D&D,L&L | | | $0_1$ |
| 55 | R1126TS444_1 | Human | nonML | D&D,D&L | | D(S),L(S) | $9_{27}$ |
| 56 | R1126TS444_4 | Human | nonML | L&L | | | $3_1$ |
| 57 | R1126TS470_1 | Human | ML | | | L(D) | $3_1$ |
| 58 | R1126TS470_2 | Human | ML | | | D(S) | $3_1$ |
| 59 | R1126TS470_3 | Human | ML | | | | $3_1$ |
| 60 | R1126TS470_4 | Human | ML | | | D(L),L(D) | $5_1\#5_2$ |
| 61 | R1126TS470_5 | Human | ML | | | D(S),L(S) | >12 |
| 62 | R1126TS489_1 | Web server | ML | D&D,D&L | | D(S),L(S) | $9_{27}$ |
| 63 | R1126TS489_2 | Web server | ML | | | D(S) | $3_1$ |
| 64 | R1126TS489_3 | Web server | ML | | | | $3_1$ |
| 65 | R1126TS489_4 | Web server | ML | | | D(L),L(D) | $5_1\#5_2$ |
| 66 | R1126TS489_5 | Web server | ML | | | D(S),L(S) | >12 |
| 67 | R1126TS490_1 | Human | nonML | | L(L) | | $0_1$ |
| 68 | R1128TS110_1 | Human | ML | D&L | | | $7_4$ |
| 69 | R1128TS238_1 | Human | ML | | | | $3_1$ |
| 70 | R1128TS238_2 | Human | ML | D&D,D&L,L&L | | L(D) | 11a121 |
| 71 | R1128TS239_3 | Web server | ML | | | L(S) | $0_1$ |
| 72 | R1128TS285_3 | Human | nonML | | | D(S) | $0_1$ |
| 73 | R1128TS416_4 | Human | ML | D&D | | | $0_1$ |
| 74 | R1128TS470_3 | Human | ML | | | L(D) | $0_1$ |
| 75 | R1128TS470_5 | Human | ML | D&L,L&L | | D(D),L(D) | $0_1$ |
| 76 | R1128TS489_3 | Web server | ML | | | L(D) | $0_1$ |
| 77 | R1128TS489_5 | Web server | ML | D&L,L&L | | D(D),L(D) | $0_1$ |
| 78 | R1136TS029_2 | Human | ML | | L(L) | | $0_1$ |
| 79 | R1136TS029_3 | Human | ML | | | | $5_2$ |
| 80 | R1136TS054_1 | Human | ML | | | | $7_2$ |
| 81 | R1136TS054_3 | Human | ML | L&L | | | 11n91 |
| 82 | R1136TS054_4 | Human | ML | | L(L) | | $0_1$ |
| 83 | R1136TS110_1 | Human | ML | D&D | | | $0_1$ |
| 84 | R1136TS110_2 | Human | ML | D&D,D&L | | L(D) | $0_1$ |
| 85 | R1136TS110_3 | Human | ML | | L(L) | L(D) | $0_1$ |
| 86 | R1136TS110_4 | Human | ML | D&D,D&L | | D(D),D(L) | $3_1$ |
| 87 | R1136TS119_2 | Human | ML | | L(L) | | $0_1$ |
| 88 | R1136TS128_4 | Human | nonML | D&L | | L(D) | $0_1$ |
| 89 | R1136TS128_5 | Human | nonML | D&L | | | $0_1$ |
| 90 | R1136TS147_3 | Human | ML | | | L(D) | $0_1$ |
| 91 | R1136TS177_1 | Human | ML | | | L(D),L(S) | $0_1$ |
| 92 | R1136TS185_1 | Human | ML | | | | $3_1$ |
| 93 | R1136TS185_2 | Human | ML | | | | $8_1$ |
| 94 | R1136TS185_3 | Human | ML | | | | $3_1$ |
| 95 | R1136TS238_1 | Human | ML | | | | TTC |
| 96 | R1136TS238_2 | Human | ML | | | | TTC |
| 97 | R1136TS238_3 | Human | ML | | | | TTC |
| 98 | R1136TS239_3 | Web server | ML | D&L | | L(L) | $8_1$ |
| 99 | R1136TS392_1 | Human | nonML | | L(L) | | $0_1$ |

| No | Model | Human/ Web server | ML/ nonML | Interlaces | Lassos | | Knots |
|---|---|---|---|---|---|---|---|
| | | | | | Shallow | Deep (>5) | |
| 100 | R1136TS416_3 | Human | ML | D&L | | L(L) | $8_1$ |
| 101 | R1136TS434_4 | Human | nonML | | | | $3_1$ |
| 102 | R1136TS444_3 | Human | nonML | | | | $5_2$#$3_1$ |
| 103 | R1136TS470_1 | Human | ML | | | | $5_2$#$3_1$ |
| 104 | R1136TS470_2 | Human | ML | D&D,L&L | | | |
| 105 | R1136TS470_3 | Human | ML | | | | $4_1$ |
| 106 | R1136TS470_4 | Human | ML | | | | $3_1$ |
| 107 | R1136TS470_5 | Human | ML | D&L | | | $4_1$ |
| 108 | R1136TS489_1 | Web server | ML | | | | $5_2$#$3_1$ |
| 109 | R1136TS489_2 | Web server | ML | D&D,L&L | | | >12 |
| 110 | R1136TS489_3 | Web server | ML | | | | $4_1$ |
| 111 | R1136TS489_4 | Web server | ML | | | | $3_1$ |
| 112 | R1136TS489_5 | Web server | ML | D&L | | | $4_1$ |
| 113 | R1138TS035_4 | Web server | nonML | D&D | | | $0_1$ |
| 114 | R1138TS054_1 | Human | ML | | | | $3_1$ |
| 115 | R1138TS054_3 | Human | ML | L&L | | L(D) | $7_2$ |
| 116 | R1138TS076_2 | Human | ML | D&D,D&L | | L(D) | $0_1$ |
| 117 | R1138TS076_3 | Human | ML | L&L | | | $0_1$ |
| 118 | R1138TS076_4 | Human | ML | D&L | | | $3_1$ |
| 119 | R1138TS076_5 | Human | ML | L&L | | | $5_2$ |
| 120 | R1138TS125_1 | Web server | ML | D&D,D&L | | L(D) | $0_1$ |
| 121 | R1138TS147_1 | Human | ML | | | | $3_1$ |
| 122 | R1138TS147_4 | Human | ML | | | | $3_1$ |
| 123 | R1138TS185_1 | Human | ML | | | L(S) | |
| 124 | R1138TS185_1 | Human | ML | | | L(S) | >12 |
| 125 | R1138TS185_3 | Human | ML | | | | $3_1$ |
| 126 | R1138TS185_4 | Human | ML | | | D(S) | $3_1$ |
| 127 | R1138TS185_5 | Human | ML | | | | $3_1$ |
| 128 | R1138TS227_2 | Human | nonML | | | | $3_1$ |
| 129 | R1138TS229_1 | Web server | ML | | | | TTC |
| 130 | R1138TS229_2 | Web server | ML | | | | TTC |
| 131 | R1138TS229_4 | Web server | ML | L&L | | | $5_2$ |
| 132 | R1138TS235_1 | Human | nonML | D&D,D&L,L&L | L(L) | D(D),D(L),L(D) | 12a1282 |
| 133 | R1138TS235_2 | Human | nonML | D&D,D&L,L&L | L(L) | D(D),L(D),L(L) | >12 |
| 134 | R1138TS238_1 | Human | ML | | | | TTC |
| 135 | R1138TS238_2 | Human | ML | | | | TTC |
| 136 | R1138TS238_3 | Human | ML | | | | TTC |
| 137 | R1138TS238_4 | Human | ML | | | | TTC |
| 138 | R1138TS239_1 | Web server | ML | | | | TTC |
| 139 | R1138TS239_4 | Web server | ML | L&L | | | $5_2$ |
| 140 | R1138TS239_5 | Web server | ML | D&D,D&L,L&L | | D(D),D(L),D(S),L(D),L(L) | $0_1$ |
| 141 | R1138TS248_4 | Human | nonML | D&D | | | $0_1$ |
| 142 | R1138TS416_1 | Human | ML | D&D,D&L | | L(D) | $0_1$ |
| 143 | R1138TS416_4 | Human | ML | L&L | | | $5_2$ |
| 144 | R1138TS434_4 | Human | nonML | | | | $3_1$ |
| 145 | R1138TS439_3 | Human | ML | D&D,D&L | | L(D) | $0_1$ |
| 146 | R1138TS439_4 | Human | ML | L&L | | | $5_2$ |
| 147 | R1138TS470_1 | Human | ML | D&D | | D(S),L(D),L(S) | TTC |
| 148 | R1138TS470_2 | Human | ML | | | D(S),L(D),L(S) | TTC |
| 149 | R1138TS470_3 | Human | ML | D&L,L&L | L(S) | D(D),D(S),L(D),L(S) | TTC |
| 150 | R1138TS470_4 | Human | ML | | | | TTC |

| No | Model | Human/ Web server | ML/ nonML | Interlaces | Lassos | | Knots |
|---|---|---|---|---|---|---|---|
| | | | | | Shallow | Deep (>5) | |
| 151 | R1138TS470_5 | Human | ML | D&L | L(S) | D(S),L(D),L(S) | TTC |
| 152 | R1138TS489_1 | Web server | ML | D&D,D&L,L&L | | D(D),D(S),L(D),L(S) | TTC |
| 153 | R1138TS489_2 | Web server | ML | | | D(S),L(D),L(S) | TTC |
| 154 | R1138TS489_3 | Web server | ML | D&D,D&L | | D(S),L(D),L(S) | TTC |
| 155 | R1138TS489_4 | Web server | ML | D&D | L(S) | D(S),L(D),L(S) | |
| 156 | R1138TS489_5 | Web server | ML | | | | TTC |
| 157 | R1149TS097_1 | Human | ML | | | L(S) | $0_1$ |
| 158 | R1149TS177_1 | Human | ML | | L(S) | | $0_1$ |
| 159 | R1156TS054_5 | Human | ML | | | L(S) | $0_1$ |
| 160 | R1156TS119_5 | Human | ML | | | L(S) | $0_1$ |
| 161 | R1156TS177_1 | Human | ML | | L(L) | | $0_1$ |
| 162 | R1156TS235_5 | Human | nonML | | | L(D),L(S) | $0_1$ |
| 163 | R1189TS238_1 | Human | ML | D&D | L(S) | D(S),L(D) | $0_1$ |
| 164 | R1190TS185_2 | Human | ML | | | L(S) | $0_1$ |
| 165 | R1190TS235_3 | Human | nonML | | L(L) | L(D) | $0_1$ |
| 166 | R1190TS238_2 | Human | ML | D&D | L(S) | D(S),L(D) | $0_1$ |
